# Quantitative prediction of ensemble dynamics, shapes and contact propensities of intrinsically disordered proteins

**DOI:** 10.1101/2022.03.21.485081

**Authors:** Lei Yu, Rafael Brüschweiler

## Abstract

Intrinsically disordered proteins (IDPs) are highly dynamic systems that play an important role in cell signaling processes and their misfunction often causes human disease. Proper understanding of IDP function not only requires the realistic characterization of their three-dimensional conformational ensembles at atomic-level resolution but also of the time scales of interconversion between their conformational substates. Large sets of experimental data are often used in combination with molecular modeling to restrain or bias models to improve agreement with experiment. It is shown here for the N-terminal transactivation domain of p53 (p53TAD) and Pup how the latest advancements in molecular dynamics (MD) simulations methodology produces native conformational ensembles by combining replica exchange with series of microsecond MD simulations. They closely reproduce experimental data at the global conformational ensemble level, in terms of the distribution properties of the radius of gyration tensor, and at the local level, in terms of NMR properties including ^15^N spin relaxation, without the need for reweighting. The IDP ensembles were analyzed by graph theory to identify dominant inter-residue contact clusters and characteristic amino-acid contact propensities. These findings indicate that modern MD force fields with residue-specific backbone potentials can produce highly realistic IDP ensembles sampling a hierarchy of nano- and picosecond time scales providing new insights into their biological function.

**AUTHOR SUMMARY:** Accurate prediction of the conformational ensemble dynamics sans bias is shown for intrinsically disordered proteins including the transactivation domain of p53.

## INTRODUCTION

Intrinsically disordered proteins (IDPs) and protein regions (IDRs) are an integral part of the proteomes of many different organisms with more than 30% of all eukaryotic proteins possessing 40 or more consecutive disordered residues.(1, 2) While IDPs and IDRs in isolation do not adopt well-defined three-dimensional (3D) structures, they often play important biological roles in molecular recognition processes by interacting in specific ways with binding partners that are typically well-ordered.(3–5) For instance, the human oncoprotein protein p53 possesses the N-terminal transactivation domain (p53TAD) that binds to the N-terminal domain of human MDM2 protein adopting a stable α-helix.(6) Prokaryotic ubiquitin-like protein (Pup) is another IDP that is directly linked to protein degradation folding into an a-helix when binding to Mpa protein.(7) In addition to binding to their target protein(s), IDPs can also be involved in liquid-liquid phase separation (LLPS).(8–11) LLPS is the segregation of molecules in solution into a condensed phase and a dilute phase with high and low biomolecular concentrations. These membraneless droplet-like compartments formed by IDPs and other biomolecules are important for cellular function. Knowledge of the structural and dynamic propensities of IDPs both in isolation and in complex biological environments is essential for understanding these processes and their role in human diseases.

In order to relate IDP sequences to biological function, detailed knowledge of IDP conformational ensembles is needed. The description of conformational ensembles can range from local secondary structure populations to explicit ensembles in 3D space with atomic resolution.(12) Some of the earliest approaches generate random coil conformational ensembles that are subsequently refined against a host of experimental data reflecting both local and global structural features.(13–15) These approaches continue to be successfully applied through integrative modeling provided that a large amount of high quality experimental data is available for each system under investigation.(16, 17) Even when data from various complementary experimental techniques are being used, the amount of experimental information obtainable is still sparse when compared to the information needed to uniquely characterize large, highly heterogeneous structural ensembles that are the hallmark of IDPs. As a consequence, the amount of information that can be gained and that is not directly reflected in the experimental data used to refine the ensemble is restricted to robust descriptors ranging from coarse-grained to global that can be compared with predictions by polymer theory under various assumptions.(16) In addition, site-specific interaction information, such as transient inter-residue contacts, can be obtained at medium to low resolution from paramagnetic relaxation enhancement (PRE) experiments by attaching electron spin labels to selected sites.(15, 17) Because empirical ensembles generated based on such data lack a time axis, they do not include dynamics time scales of IDPs associated with interconversion rates between substates and, hence, they do not inform about an essential part of the energy landscape.

From a theoretical and computational perspective, all-atom molecular dynamics (MD) simulations are an attractive alternative to empirical approaches for the generation of IDP conformational ensembles, including dynamic time scale information, for the comprehensive interpretation of experimental results.(18) However, for many years limitations in computer power precluded the generation of statistically well-converged results and MD force fields primarily developed for ordered proteins turned out to be unsatisfactory for applications to IDPs. With the continuing increase in computer power, the quality of sampling has reached a level that allows rigorous validation by quantitative comparison with a rich body of experimental data. In cases where discrepancies are observed between simulation and experiment, as is commonly the case, approaches have been developed that use restraining or reweighting that bias the original simulation to obtain results that agree better with experimental data.(19–26) When not only the conformational ensemble but also the underlying dynamics time scales are of interest, suitable rescaling of the MD time step or correlation times of the dominant motional modes can be applied to improve agreement with experiment.(27–30) Because these methods can often improve the unaltered simulations only within certain boundaries, they are best suited when the original predictions are fairly close to experimental data.(31) Although these methods rarely fail to produce better agreement, at least on average for those experimental parameters directly used as restraints or for reweighting, they naturally depend on large amounts of experimental data of good quality as input for each protein system studied. This amounts to a laborious experimental effort that needs to be repeated for each new protein system as the experimental data are protein-specific rendering them non-transferrable between systems.

An alternative and more principled approach is to improve the MD force fields themselves enabling them to increasingly accurately predict experimental data in a way that is fully transferrable between protein systems, both ordered and disordered. This premise has led to a recent proliferation of protein force field developments(32–37) and new explicit water models(38–40) specifically geared toward the improved representation of disordered proteins. In a significant development, residue-specific force fields have been introduced.(41) These force fields use in addition coil library information from the Protein Data Bank (PDB) by incorporating the individual backbone φ,ψ propensities of each residue type.(41–47) Such residue-specific force fields, in combination with suitable water models, can provide an improved representation of disordered states while retaining the properties of ordered proteins. With respect to water models, TIP4P-D and closely related derivatives have been notably successful in preventing overly compact conformations by favoring more extended IDP structures showing improved agreement with experiment. (38)

Besides global properties, such as the radii of gyration and asphericities, IDP ensembles and trajectories should also accurately reproduce local dihedral angle distributions and secondary structure propensities. Moreover, they should also replicate dynamic and kinetic IDP properties, such as librational motions and time scales of interconversion between conformational substates. Such information is important for understanding recognition events between IDPs and their binding targets, including IDP interactions with other disordered biomolecules, for example, during the formation of LLPS condensates. Experimental IDP dynamics information can be gained from fluorescence depolarization spectroscopy,(48) Förster resonance energy transfer (FRET),(16) and nuclear magnetic resonance (NMR) relaxation.(15) NMR ^15^N longitudinal *R*_1_ and transverse *R*_2_ spin relaxation rates are exquisitely sensitive to the dynamics of disordered proteins and the underlying time scales.(49–51) *R*_2_ relaxation rates, for example, have been linked to residual intramolecular interactions in chemically unfolded proteins.(51–53) ^15^N *R*_1_ and *R*_2_ rates can be experimentally determined for each protein residue and therefore they are valuable for validating MD simulations with respect to amplitudes and time scales of IDP dynamics.(29, 54–56)

We recently developed the AMBER ff99SBnmr2 force field by modifying the backbone dihedral angle potentials of each amino-acid residue type to reproduce the φ,ψ dihedral angle distributions found in a random coil library.(57) The ff99SBnmr2 force field has been validated against experimental nuclear magnetic resonance (NMR) scalar ^3^*j*-couplings of α-synuclein and β-amyloid IDPs demonstrating that this force field accurately reproduces their sequence-dependent local backbone structural propensities.(58) The primary goal of this work is to learn whether state-of-the-art replica exchange and extended MD simulations of IDPs can also realistically reproduce NMR *R*_1_, *R*_2_ relaxation rates with their strong and unique dependence on motional time scales without the need of any additional corrections such as constraints or reweighting. Moreover, in-depth analysis of the MD trajectories generated yields a wealth of information about the radius of gyration tensor distribution and dominant dynamics modes allowing graph-theory based identification of specific inter-residue interaction propensities and residue clusters for the better understanding of IDP behavior.

## RESULTS

### Ensemble properties of radius of gyration tensor

The radius of gyration *R*_g_(t) is shown as a function of time for representative 1-μs MD trajectories of p53TAD and Pup in **Fig. 1A,B** (see also **Fig. S1**). The trajectories exhibit predominantly stationary stochastic behavior reflecting random expansion and contraction of the overall IDP size with the mean value (blue horizontal lines) in good agreement with the experimentally determined <*R*_g_> (black line) or the predicted <*R*_g_> from polymer theory (**Eq. 6**). The MD-distributions of *R*_g_ of all 10 MD trajectories are shown as histograms in **Fig. 1C,D**. The Flory exponent ν of the polymer scaling law was determined from the REMD ensembles at 298 K. Using ρ_0_ = 1.927 Å, we obtain a value of ν = 0.601 for Pup, which closely matches the theoretical value ν_theory_ = 0.588 of a fully disordered, self-avoiding random coil.(59, 60) For p53TAD, the <*R*_g_> value of 28.1 Å is in almost perfect agreement with experiment (28.0 Å) corresponding to ν = 0.624, which clearly exceeds ν_theory_.

**Fig. 1.**
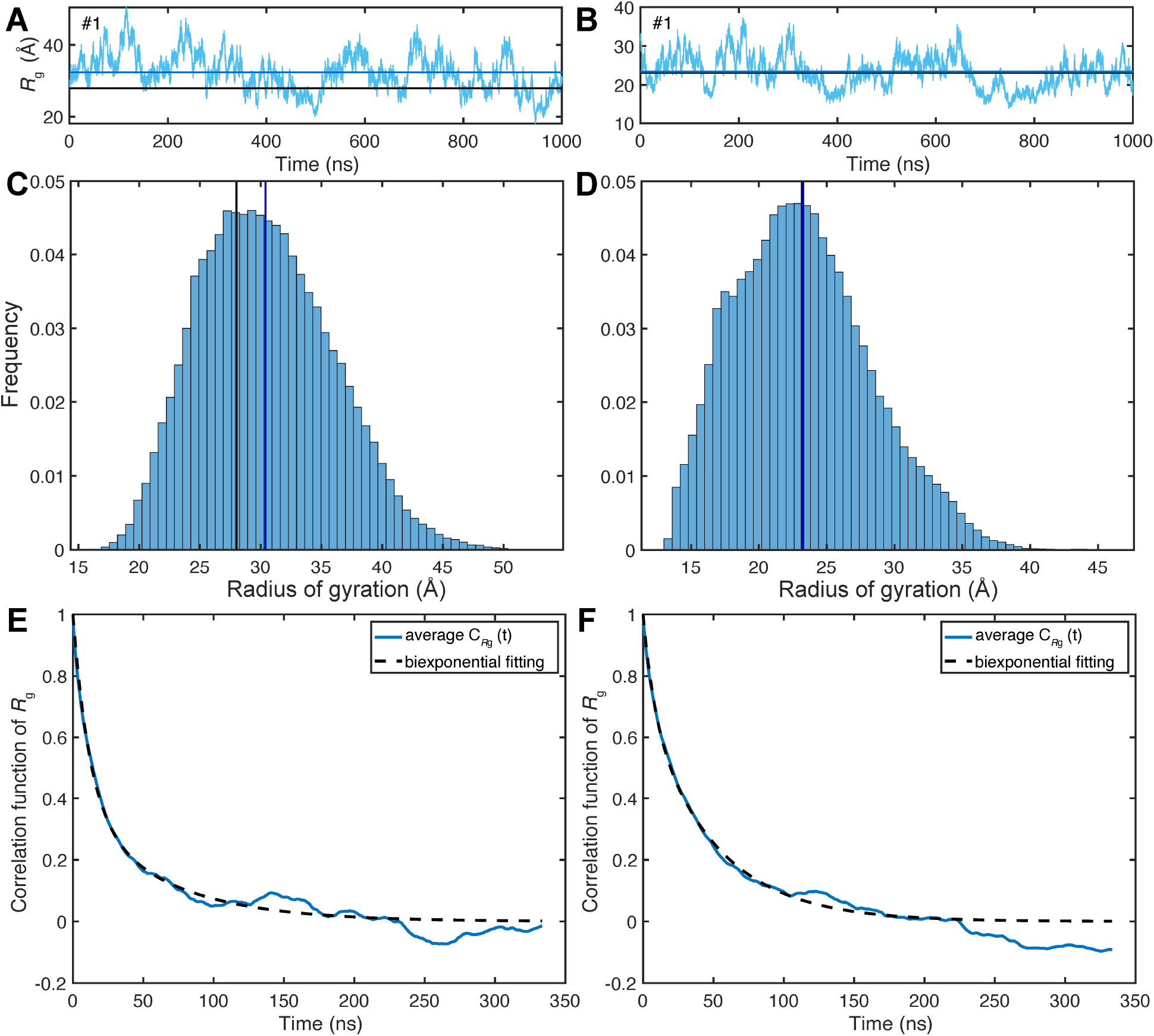
Radius of gyration, *R*_g_, properties of two IDPs p53TAD and Pup from microsecond MD simulations. Time-dependence of *R*_g_(t) from representative 1-μs MD trajectories (cyan) of (A) p53TAD and (B) Pup where the horizontal blue lines correspond to the mean *R*_g_ values calculated from the trajectories and the black lines correspond to the experimentally determined *R*_g_ for p53TAD and the predicted *R*_g_ according to polymer theory for Pup. *R*_g_ profiles for all 10 1-μs trajectories of each protein are shown in Figure S1. Histograms of the *R*_g_(t) distributions over all 10 MD simulations are shown in Panels C, D (blue and black lines have the same meaning as in Panels A, B). The standard deviation of *R*_g_ over all 10 MD trajectories is 5.4 Å for p53TAD and 5.0 Å for Pup. Offset-free time-correlation functions *C*_*R*g_(t) of *R*_g_(t) averaged over all 10 1-μs MD trajectories are shown for (E) p53TAD and (F) Pup. The dashed lines belong to non-linear least squares fits of *C*_*R*g_(t) by biexponential functions whereby the best fits are obtained for p53TAD with τ_a_ = 12 ns (63% of total amplitude), τ_b_ = 62 ns (37%) and for Pup with τ_a_ = 8 ns (29%), τ_b_ = 48 ns (71%).

The characteristic time scales of *R*_g_(t) fluctuations can be obtained from the time-correlation functions *C*_*R*g_(t) (**Eq. 5**), which are well-converged over the course of the 1-μs trajectories (**Fig. 1E,F**). *C*_*R*g_(t) of both proteins decay in good approximation biexponentially with reconfigurational correlation times τ_a_ ≅ 10 ns and τ_b_ ≅ 55 ns. The normalized variance of the *R*_g_(t) fluctuations, given by

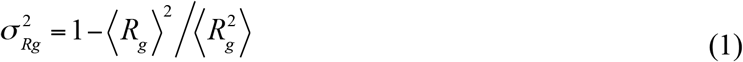

is almost the same for p53TAD (0.03) and Pup (0.04). The ensemble distribution of the gyration tensor ***S*** (**Eq. 2**) contains information about the deviation of individual MD snapshots from spherical shape, which can be directly compared with a random Gaussian chain serving as a perfect random coil (**Fig. 2**).(61) Both proteins show unimodal asphericity distributions (**Eq. 3**) with maxima around *A* ≅ 0.18, which qualitatively differ from the Gaussian chain model (**Fig. 2C**) peaking at *A* = 0. Compared to p53TAD, Pup has a higher tendency to adopt a more spherical conformation. Another useful measure of the overall shape of individual snapshots is the prolateness *P* (**Eq. 4**). The distribution of *P* is bimodal for both proteins with the global maximum corresponding to prolate-shaped (cigar-like) structures (*P* = 1) and a second (local) maximum corresponding to disk-like structures (*P* = −1). The distribution of the prolateness of Pup is more balanced between positive and negative values with <*P*> = 0.2 than for p53TAD, which has a higher tendency to adopt prolate-shaped conformers (<*P*> = 0.35), whereas the Gaussian chain distribution (<*P*> = 0.3) lies between the two IDP distributions. The distinct asphericity distribution and increased prolateness of p53TAD is at the origin of its increased <*R*_g_> over the Gaussian random coil model.

**Fig. 2.**
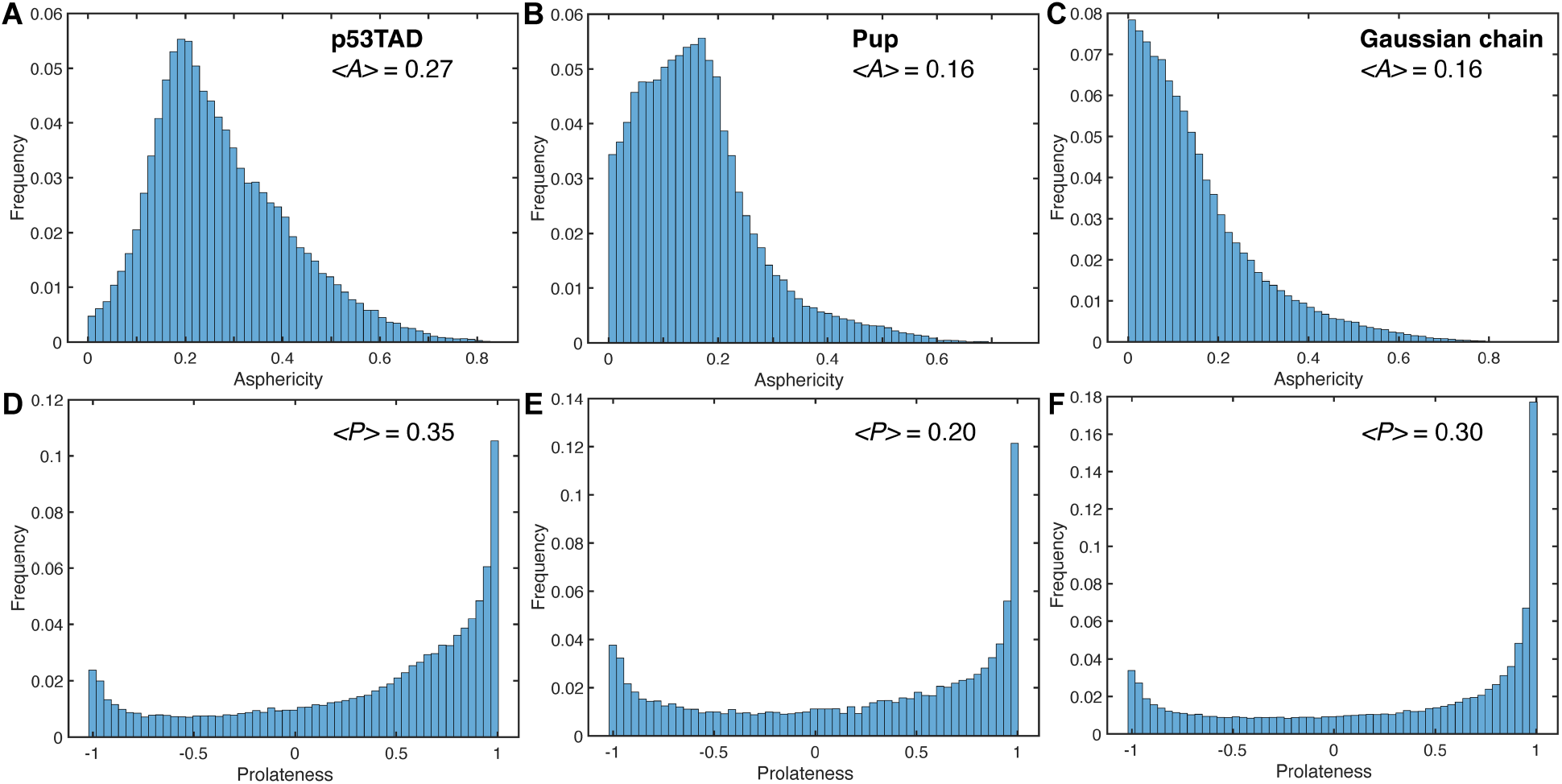
Gyration tensor properties of IDP ensembles of p53TAD and Pup across 10 1-μs MD trajectories. The distributions of gyration tensor aspherities *A* are shown for (A) p53TAD and (B) Pup in comparison with (C) a Gaussian chain. The distributions of gyration tensor prolateness *P* are shown for (D) p53TAD and (E) Pup in comparison with a (F) Gaussian chain.

### Validation against *R*_1_, *R*_2_ relaxation data

Experimental and computed ^15^N *R*_1_, *R*_2_ relaxation rates are shown in **Fig. 3**. *R*_1_ relaxation rates determined from simulations (**Eq. 7–12**) are in close agreement with experiment evidenced by small RMSEs (0.10 s^-1^ for p53TAD and 0.12 s^-1^ for Pup) and Pearson correlation coefficients R of 0.78 for p53TAD and 0.86 for Pup (**Fig. 3A,B**). *R*_2_ relaxation rates determined from the simulations are also in good agreement with experiment with correlation coefficients R of 0.88 for p53TAD and 0.70 for Pup and RMSEs of 0.84 s^-1^ for p53TAD and 0.81 s^-1^ for Pup and (**Fig. 3C,D)**. It can be seen that the simulations tend to underestimate *R*_1_ and overestimate *R*_2_ rates, although only slightly, in a manner that is notably uniform for the *R*_1_ values of both proteins and for the *R*_2_ values of p53TAD. The 10 N-terminal residues of p53TAD are very flexible with small *R*_2_’s, which closely follow the experiment. For Pup, differences in *R*_2_ between MD and experiment display the same trend and are most pronounced for residues 30–48. The error bars of the computed relaxation rates, which represent the root-mean-square deviations over all 10 MD trajectories, are fairly uniform along the polypeptide chains and systematically larger for *R*_2_ than for *R*_1_, again with the exception of the 10 N-terminal residues of p53TAD. For both proteins, not all 10 1-μs MD trajectories individually reproduce the experimental data equally well. Either 1 (p53TAD) or 2 (Pup) trajectories have more compact average IDP structures, which quantitatively affect the agreement with experiment (**Fig. S2**).

**Fig. 3.**
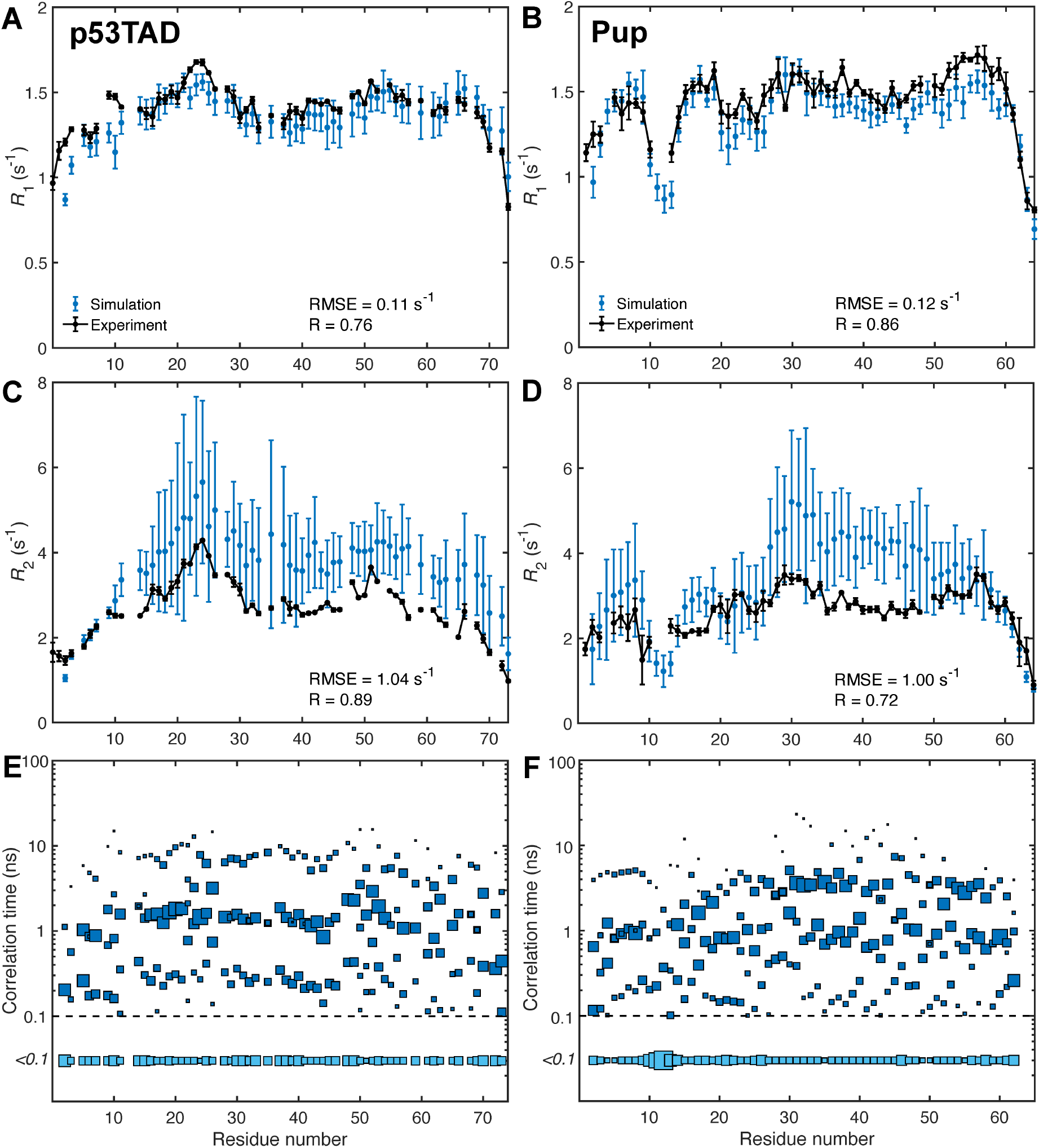
Back-calculated *R*_1_, *R*_2_ NMR ^15^N-spin relaxation rates in comparison with experiment along with underlying motional time scale distributions. *R*_1_, *R*_2_ rates calculated from average correlation functions are plotted in blue with error bars representing standard deviations across individual MD trajectories. Correlation time distribution of individual ^15^N-^1^H bonds of IDPs extracted from correlation functions for (E) p53TAD and (F) Pup where the size of the blue squares are proportional to the associated motional amplitudes *A*_i_. The squares at the bottom indicate the aggregate of dynamics contributions with correlation times faster than 100 ps. Dominant dynamics time scales range from about 100 ps to about 10 ns depending on the residue, with the exception of Thr12 in Pup which exhibits dominant dynamics time scales faster than 100 ps.

Correlation times of backbone N-H bond vectors in both proteins fitted from the average correlation functions range from picoseconds to about 20 ns (**Fig. 3E,F**). Consistent with the finding for other IDPs,(55, 62) the dominant contribution to the time correlation functions stems from dynamics on the intermediate time scale around 1 ns reporting about backbone φ,ψ jumps. Fast dynamics on the time scale of 100 ps or faster report on local ^15^N-^1^H bond librations, similar to those observed in secondary structures of folded proteins,(63) and slower dynamics on the time scale between 3 and 20 ns reports on collective IDP chain motions. The presence of slower modes correlate with increased *R*_2_ values most pronounced for residues 30–48 in Pup. This is consistent with relaxation theory (**Eq. 12**), which predicts that in solution transverse spin relaxation rates *R*_2_ are in good approximation proportional to the effective overall correlation time experienced by the ^15^N-^1^H spin pairs.

Increased transverse NMR spin relaxation is indicative of the presence of collective segmental motions in IDPs, which are modulated by the formation of transient secondary structures and inter-residue side-chain interactions. To examine these relationships, instantaneous secondary structures and average contact maps were determined from the MD trajectories (**Fig. 4**). A contact is defined in an MD snapshot when the nearest distance between atoms from two different residues is smaller than 4 Å (uninformative first-neighbor (*i*,*i*+1) and second-neighbor (*i*,*i*+2) contacts between residues were excluded (white band along diagonal in **Fig. 4A,B**)). The most frequent contacts are relatively short range, but contacts over larger distances occur for p53TAD and even more frequently for Pup. Some contacts are linked to the transient formation of short secondary structures, α-helices and β-strands (**Fig. 4C,D**), whereas other regions display frequent contacts largely independent of secondary structure propensity often involving arginine residues, such as Arg65 of p53TAD and Arg28/29 and Arg56 of Pup. **Fig. 4C,D** also shows that selected trajectories possess regions with well above-average secondary structure propensities, such as trajectories #4 of p53TAD and trajectories #5 and #7 of Pup, which are the same trajectories that contribute to the lengthening of *R*_2_ along parts of the polypeptide sequences mentioned above. Due to their atypical nature, not representative of the other trajectories, they were excluded for some of the following residue-cluster analysis. For p53TAD, regions that tend to form α-helices do not form β-strands and vice versa (except for trajectory #4). For Pup, on the other hand, a number of regions exist in its N-terminal half that can transiently switch between these two types of local secondary structures.

**Fig. 4.**
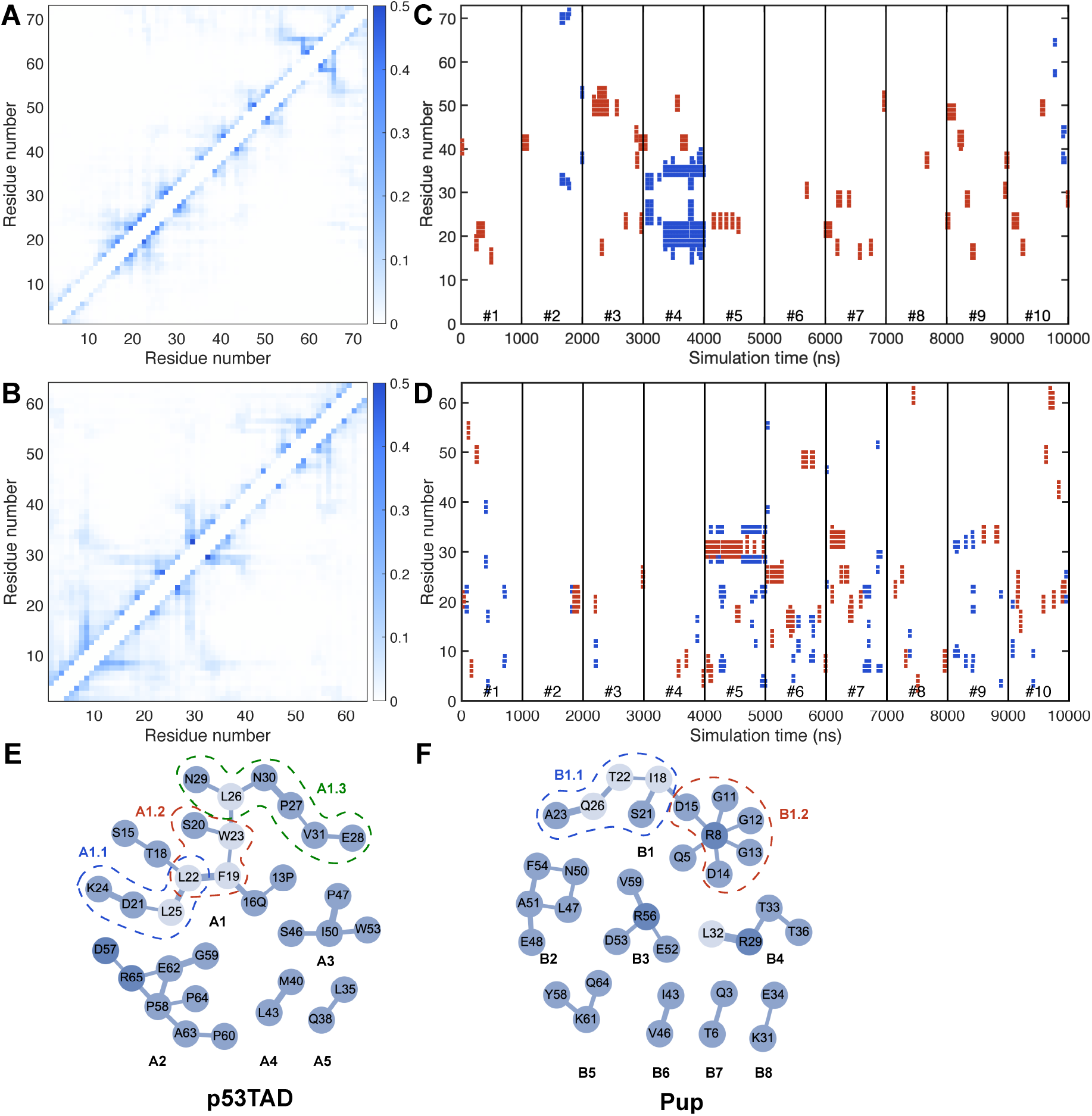
Average IDP contact maps and time-dependent secondary structure formation of each residue. (A, B) Pairwise contact occupancies were determined from MD simulations (without atypical trajectories, **Fig. S2**) for (A) p53TAD and (B) Pup. Darker/lighter shades of blue denote contacts that are more frequently/rarely formed according to legend (vertical bar). Self-contacts, first-neighbor contacts (between residues *i*,*i*+1), and second-neighbor contacts (between residues *i*,*i*+2) are not shown since they are present in most snapshots. (C, D) secondary structure of each residue in MD simulations are predicted using the DSSP algorithm with α-helices shown in red and β-strands in blue. (E, F) In the residue clusters at the bottom, pairwise contacts with occupancies > 0.2 are depicted as an edge connecting two nodes (residues) with edge widths proportional to the pairwise contact occupancies. Labels A1–A5 denote dominant clusters in p53TAD and B1–B8 in Pup. Examples of transiently formed subclusters are indicated by dashed lines (A1.1, A1.2, and A1.3 in p53TAD and B1.1 and B1.2 in Pup).

### Inter-residue contact propensities

Different residues along the polypeptide chain display different tendencies to form contacts with other residues. **Fig. 5A,B** shows the average number of contacts per snapshot for each residue, which was calculated as the total number of contacts formed by a residue divided by the total number of MD snapshots. To better visualize the different behaviors, the residues were divided into four distinct groups: the majority of residues that form 0.5–1.5 contacts per snapshot (colored in black), residues that form an unusually small number of contacts (< 0.5) (colored in blue), residues that form a moderately large number of contacts (1.5–2) (colored in yellow), and residues that form a relatively large number of contacts (> 2) are colored in red. For Pup, there are three distinct regions that form the largest numbers of contacts (red) comprising residues (1) Lys7, Arg8, (2) Arg28, Arg29, and (3) Arg56. They perfectly align with the three centers of **Fig. 3** with elevated *R*_2_ values, namely (1) Arg8, (2) Arg29, and (3) Arg56. For p53TAD, the residue that forms the largest number of contacts is Arg65, which is surrounded by residues with a number of contacts below average between 0.5 and 1.0. This rationalizes why *R*_2_ of Arg65 shows a local maximum that is still lower than *R*_2_ in other regions of p53TAD, such as residues 19–26 forming a residue cluster with an intermediate number of contacts. Notably, the 11 N-terminal residues of p53TAD display a lower-than-average amount of contacts, which is consistent with low *R*_2_ values observed across all 10 individual MD trajectories. When the same type of contact analysis is performed with side-chain atoms only, a similar behavior is observed with only a small, systematic reduction in contacts (**Fig. S3**) reflecting that the majority of medium-to long-range inter-residue contacts are made by side-chain atoms.

**Fig. 5.**
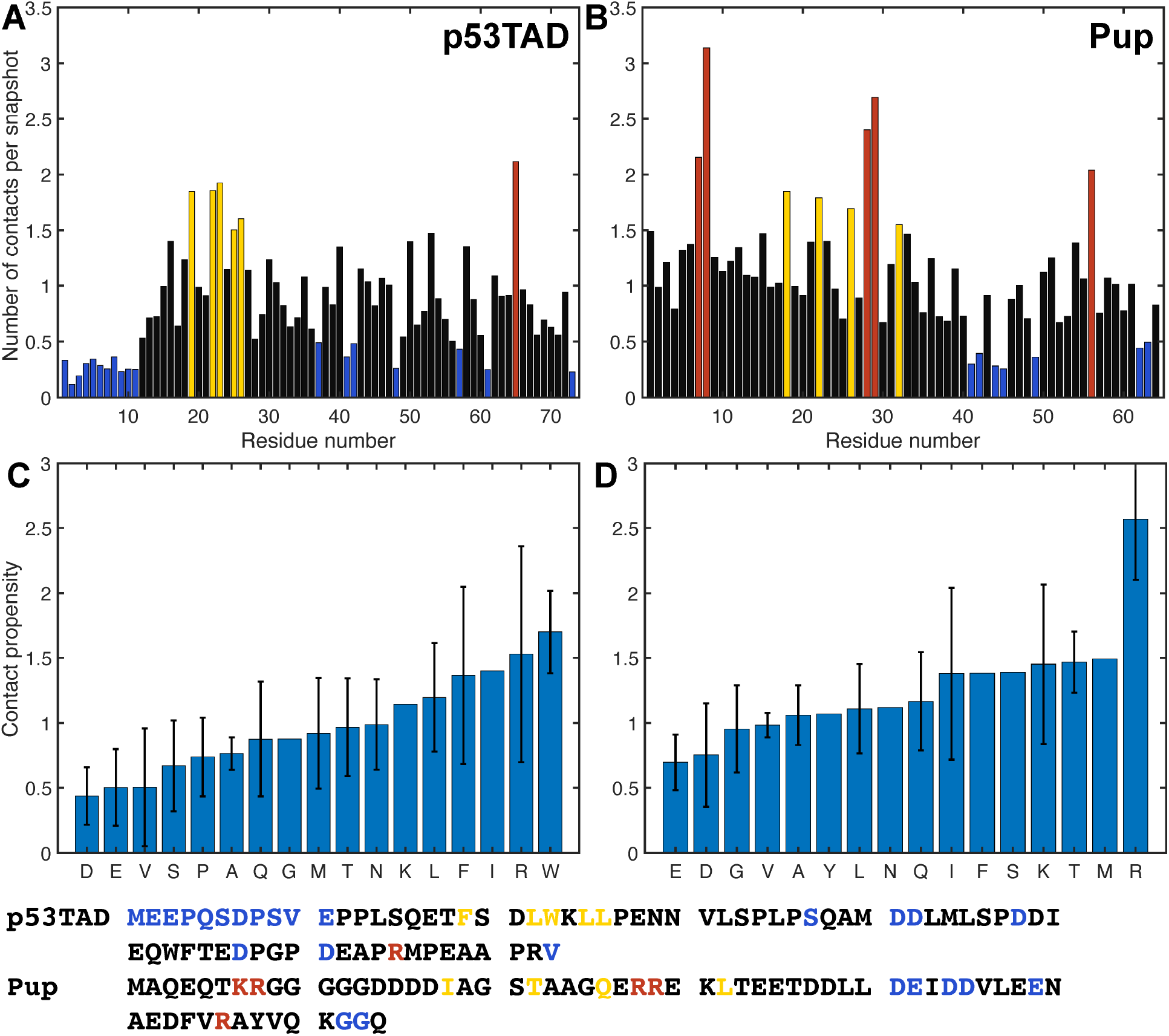
Number of close contacts formed by each residue during MD simulations of p53TAD and Pup (without outliers) along with average residue-type specific contact propensities. For each residue, the number of contacts was normalized by the number of snapshots for (A) p53TAD and (B) Pup. Residues with their number of contacts per snapshot below 0.5 are depicted in blue, 0.5–1.5 in black, 1.5–2 in yellow, and above 2 in red. Primary sequences of p53TAD and Pup are given at the bottom and colored as in Panels A, B. Average contact propensities according to amino-acid residue type, which is the number of contacts per snapshot averaged over all residues of the same type, are shown for (C) p53TAD, (D) Pup. Error bars correspond to the standard deviations among different residues of the same type.

We also grouped the number of contacts per snapshot formed by each residue according to residue type and normalized them by the number of residues of the same type. The resulting value for each amino acid residue type present in p53TAD and Pup reflects their inherent contact propensity (**Fig. 5C,D**). These profiles display the following trends: positively charged residues arginine and lysine are on average most prone to form contacts, followed by hydrophobic residues isoleucine and leucine as well as aromatic residues tryptophan and phenylalanine. Negatively charged residues aspartate and glutamate, however, are least disposed to form contacts. This may be also a consequence that both IDPs are overall negatively charged (−14e for p53TAD and −12e for Pup). When acidic residues outnumber basic residues, the former tend to repulse each other, thereby increasing *R*_g_, while the latter have more options to interact with an acidic residue than vice versa leading to an increase of the contact propensity of basic over acidic residues.

### Contact analysis by graph theory

To investigate the nature of some of the most frequent pairwise contacts in these IDPs, the MD snapshots were analyzed by graph theory where each snapshot is represented as an undirected graph with each residue corresponding to an edge and an inter-residue contact corresponds to an edge connecting the two residues (nodes). The resulting graphs were then analyzed in terms of clusters, which are disconnected graph components that do not have any edges to nodes outside of the cluster. On average 6.0 clusters per snapshot are found for p53TAD and 5.4 clusters for Pup. The probabilities of a cluster to have a given size are represented for both IDPs by the histograms of cluster sizes (**Fig. 6A**), which reveal that clusters consisting of 2 nodes are most abundantly present (around 40%) in both p53TAD and Pup. Moreover, the cluster size probability decreases rapidly with increasing size. For instance, the fraction of clusters with 10 or more nodes (residues) is only 2–3%. Despite their sequence independence and different lengths, the two IDPs have strikingly similar cluster size distributions. The number of edges grows on average linearly with the number nodes (straight solid line), which is much slower than the quadratic behavior of complete graphs (dashed line, **Fig. 6B**). In fact, most of the clusters formed during MD simulations are sparse graphs with a relatively small average edge-to-node ratio of 1.54, which is indicative of tree-like graphs consisting mostly of linear branches with few cross-links. **Fig. 6** also depicts residue clusters (on the right) where pairwise contacts with occupancies > 0.2 are depicted as an edge connecting two nodes (residues) with edge widths proportional to the pairwise contact occupancies.

**Fig. 6.**
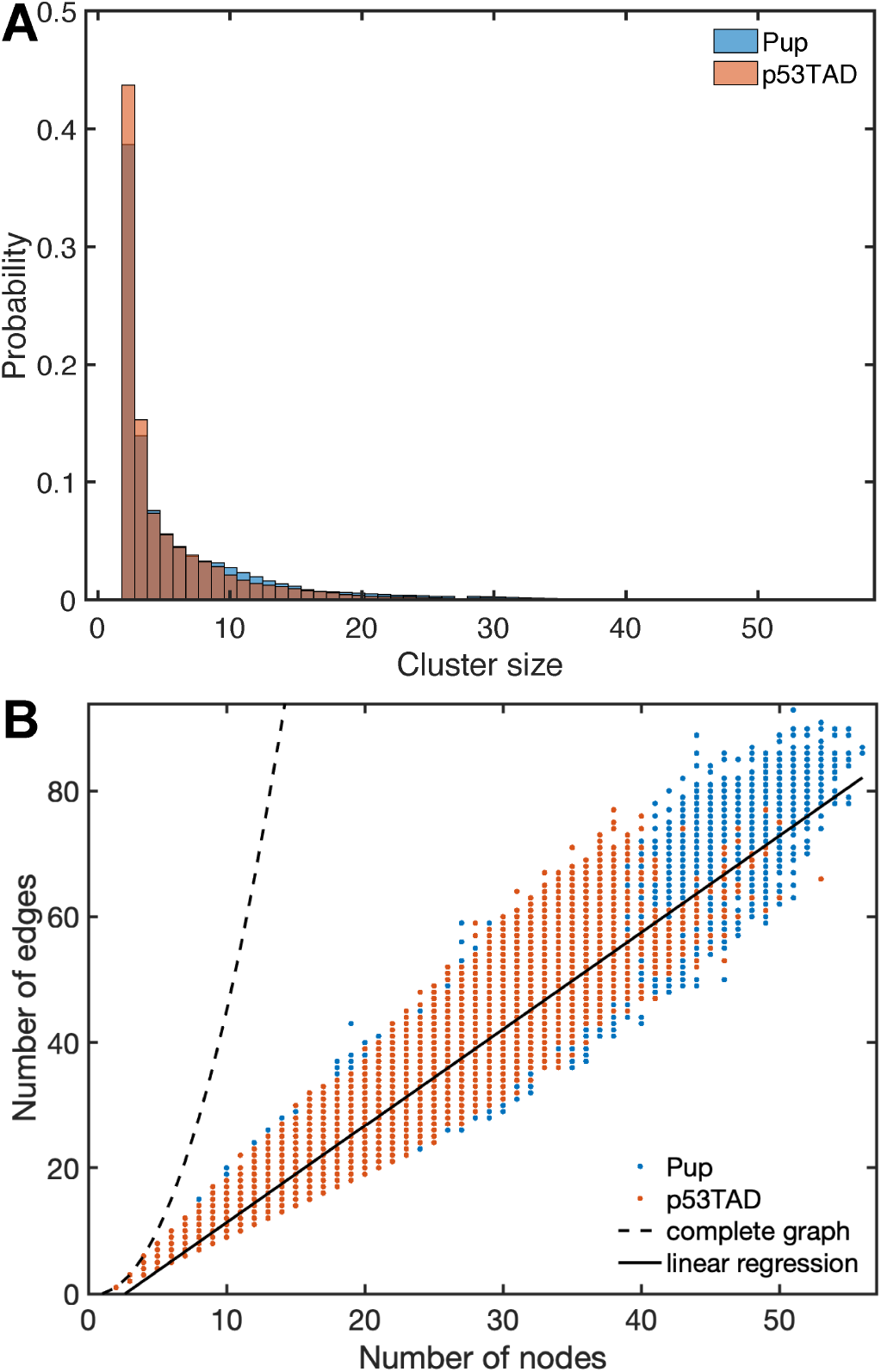
Graph theoretical analysis of inter-residual interactions and transient interaction networks of p53TAD and Pup. (A) Clusters consisting of 2 nodes (residues) dominate in the MD structures of p53TAD and Pup (without outlier trajectories), followed by clusters of size 3, etc. (B) The majority of the unique clusters are sparse graphs, with their number of edges much smaller than the number of edges in complete graphs growing with *N*(*N*-1)/2 where *N* is the number of nodes. The average edge-to-node ratio is 1.54 (slope indicated by solid black line), indicating predominantly tree-like graphs that sometimes have a few additional edges (cross-linked branches).

The graph-theoretical representation of the transient interaction network uncovers the relationship between *R*_2_ profiles and transient contact formation and the types of interactions that are prevalent in IDP structures. For p53TAD, the three centers in the sequence with an elevated experimental *R*_2_ profile are (1) Lys24, (2) Glu51, and (3) Met66, and they are involved in or are sequentially adjacent to clusters A1, A3, and A2, respectively. Electrostatic interactions are important for residue cluster formation in p53TAD, in particular in cluster A2 featuring the pairwise contacts Lys65–Asp57 and Arg65–Glu62. The largest elevation of *R*_2_, however, is the result of the largest interaction network A1. Hydrophobic and aromatic residues Phe19, Leu22, Trp23, Leu25 and Leu26 belong to a p53TAD segment that displays increased helical propensity(64, 65) (secondary structure propensities determined from chemical shifts are shown in **Fig. S5**) and which undergoes distinct loop closure dynamics. (66) In particular, residues Phe19, Trp23, and Leu26 form the hydrophobic triad that is crucial for the binding of p53TAD to MDM2.(65) Similar to cluster A1, the smaller cluster A3 centered around Ile50 is also driven by hydrophobic interactions.

The regions of Pup with elevated *R*_2_ values (**Fig. 3D**) around Arg8, Ile18, Thr22, Arg29, Arg56 are all involved in clusters B1, B4, or B3 (**Fig. 4E,F**). Separate clusters can involve sequentially adjacent residues, such as clusters B2 and B3 or clusters B3 and B5 and thereby mediate cooperative behavior. The most dominant inter-residue interaction in Pup is of electrostatic nature resulting in the transient formation of salt bridges involving residue pairs in cluster B1.2 (Arg8–Asp14, Arg8–Asp15) and cluster B3 (Arg56–Asp53, Arg56-Glu52). Many of these residues appear to play the role of hubs promoting enhanced interactions also with other residues as visualized by the graphs in **Fig. 4E,F**.

## DISCUSSION

Disordered proteins play a prominent role in many regulatory processes using their unique malleability to interact with their targets. Details of conformational substates of IDPs and how they are shaped by the complex interplay of inter-residue interaction networks are currently poorly understood both experimentally and computationally. In this work, we showed how the latest advances in MD force fields and computational protocols allow the nearly quantitative prediction of the complex behavior of the two IDPs p53TAD and Pup, including their dynamics time scales from site-resolved NMR spin relaxation.

The global dimensions of IDPs can be experimentally characterized by SAXS providing information about their radius of gyration *R*_g_ for direct comparison with MD ensembles. For Pup, <*R*_g_> from the 10 1-μs MD simulations follows the power law of **Eq. 6** with a Flory exponent ν = 0.601, which closely mirrors the behaviour of a self-avoiding random coil (ν = 0.598). By contrast, p53TAD is more expanded with ν = 0.624, which is consistent with previous experimental results reported for this protein.(67) Such behaviour could be the result of stronger repulsive intra-residual forces caused by a slightly higher negative net charge (−14e of p53TAD vs. −12e of Pup) and a high percentage of prolines (18% in p53TAD vs. none in Pup) known to increase extendedness.(68) The relatively high v values of both proteins suggest that their interactions with water solvent are highly favorable preventing the hydrophobic collapse of their polypeptide chains.

The 10 1-μs MD trajectories allow extensive sampling of the radius of gyration over time and extract characteristic time scales from its autocorrelation function (**Fig. 1**). For both proteins, the time-correlation function follows in good approximation a biexponential decay with correlation times around 10 and 55 ns. Global distance fluctuations can be studied experimentally by nanosecond fluorescence correlation spectroscopy (nsFCS), which found for 8 M urea denatured ubiquitin global reconfiguration times τ_r_ in the range of 50–90 ns.(16) A nsFCS study of α-synuclein, which is about twice as long in sequence as the IDPs studied here, identified two reconfigurational correlation times of τ_r1_ = 23 ns and τ_r2_ = 136 ns.(30) These correlation times are within a factor 2–3 of those found in the current study, although it should be kept in mind that they report about a donor/acceptor pair, i.e. S42C/T92C in the case of α-synuclein, rather than about *R*_g_.

Heteronuclear ^15^N relaxation offers a complementary view of IDP dynamics. Longitudinal *R*_1_ and transverse *R*_2_ relaxation rates are caused by local spin interactions, namely the magnetic dipole-dipole coupling and chemical shielding anisotropy, and they reflect reorientational dynamics amplitudes and timescales due to local conformational fluctuations as well as longer-range reorientational motional modes of the order of an IDP’s persistence length and beyond. Model-free analysis is not applicable to IDP relaxation data due to the absence of a well-conserved global rotational diffusion tensor as reference frame.(27) Instead, a residue-by-residue interpretation can applied where the correlation function of each site is described as a multiexponential function of the type of **Eq. 8** with 6 exponential dynamics modes.(28, 50, 55, 62, 69) The hierarchy of dynamics modes depicted in **Fig. 3** shows a broad distribution of time scales including rapid librational motions (< 100 ps) and dominant low nanosecond motions, which sample the different local energy basins of backbone φ,ψ dihedral angles. The slowest modes with time scales in the range of 3–20 ns represent predominantly collective segmental reorientational motions. A similar hierarchy of time scales has been observed by fluorescence depolarization kinetics measurements of α-synuclein.(48) These collective motions involve medium to longer-range interactions between residues that can be elucidated by graph theoretical analysis of the MD trajectories described here. For Pup, many of these slower motional modes have correlation times around 3–4 ns whereas for p53TAD they are on average twice as large. For both proteins the three distinct bands of time scales are pervasive across their polypeptide sequence (**Fig. 3E,F**).

MD methodology has made great strides in recent years to toward an increasingly realistic representation of disordered proteins.(26) Besides experimental scattering data, quantitative NMR has played a key role for the independent validation of MD ensembles. Because NMR spin relaxation parameters fully quantitatively reflect IDP dynamics at atomic-level resolution both in terms of motional amplitudes and time scales, their accurate reproduction by MD has been an important but also very challenging task. A recent comparison of commonly used MD force fields that do not use residue-specific backbone potentials showed for several IDPs significant force-field dependences with the best results obtained when the analysis was restricted to average correlation functions of chunks of 10-ns subtrajectories.(56) The need to exclude slower time-scale motions, which are prominent in both experimental data and simulations (see for example **Fig. 3**), may reflect the lack of convergence due to limited sampling. Beneficial for all simulations was the improvement of the TIP4P-D water model over TIP3P preventing overly collapsed IDP ensembles, which is consistent with other computational studies.(38, 57) Because of the observed discrepancies between experiments and MD simulations, some studies applied *post factum* adjustments to the MD simulations in order to improve agreement, which include uniform or selective scaling of the MD time scale or correlation times(27–30) or the reweighting of sub-trajectories.(62) Here, we chose a different approach: rather than relying on *post factum* modifications, we use the residue-specific ff99SBnmr2 force field, which was specifically designed for the improved representation of IDPs without the need of any corrections.(57, 58) A correction-free MD approach has recently been reported for the intrinsically disordered SH4UD protein with the Amber ff03ws force field, which does not use residue-type independent backbone dihedral angle potentials, and no time-scale dependent data, such as NMR spin relaxation, were used for validation.(70) NMR chemical shifts were back-calculated using SHIFTX2,(71) which, besides 3D structural information, makes extensive use of protein sequence data. Here, we back-calculated NMR chemical shifts using PPM(72) (**Fig. S4**), which only uses the physical parametrization of chemical shifts with respect to 3D protein structure of each snapshot,(71) achieving very good agreement.

The close correspondence observed between experimental and computed ^15^N relaxation *R*_1_ and *R*_2_ relaxation rates for both IDPs studied here (**Fig. 3**), without the need for *post factum* corrections, attests to the accuracy and robustness of the computational protocol used. It applies REMD for the generation of conformational ensembles belonging to different temperatures from which 10 representative structures at 300 K were randomly selected as starting structures for 1-μs MD trajectories whereby all simulations made use of the ff99SBnmr2 force field and the TIP4P-D water model. MD-derived longitudinal ^15^N *R*_1_ follow the shapes of the experimental *R*_1_ profiles with a small tendency to underestimate the experimental ^15^N *R*_1_ rates by 4–6% whereas ^15^N *R*_2_ relaxation rates overestimate the experimental values on average by 26% for Pup and 34% for p53TAD. This level of agreement is significantly better than for previously reported comparisons of this type. It is possible to achieve additional improvement by removing 1–2 MD trajectories starting from the most compact initial structures, a strategy proposed in the ABSURD method (**Fig. S3**). Although *post factum* modifications can provide better agreement with experiment, it is generally not obvious whether the altered ensembles are in fact consistent with a modified, physics-based force field. If such a connection can be established it will allow, in principle, the further improvement of force fields for applications also to other proteins. Indeed, the ff99SBnmr1 force field, which is the parent force field of ff99SBnmr2, was developed and optimized using this strategy by the systematic reweighting of MD snapshots based on many trial force fields using experimental NMR data of intact proteins.(73)

The good agreement of the MD simulation with experimental observables both motivates and justifies the analysis of other protein properties observed in the MD trajectories that are difficult to measure. This includes the analysis of transient inter-residue interactions. The molecular driving forces of these interactions are fundamentally similar to those of ordered proteins although average hydration properties may differ.(70) In contrast to ordered proteins, inter-residue interactions between non-sequential amino acids are short-lived. Therefore, the time-averaged interaction maps (**Fig. 4A,B**) offer only partial insights as they conceal the compositions and distributions of instantaneous interaction clusters. In fact, the relatively large network reflected by the average contact map contrasts the much smaller size of graphs that exist at any given time, which attests to the very heterogeneous and transient nature of instantaneous contact clusters. The highest occupancy of pairwise contacts found is around 0.5, which mostly belong to (*i*,*i*+3) contacts. For a list of the most frequent pairwise contacts, **see Tables S2, S3**.

Snapshot by snapshot analysis revealed the dominance of small cluster sizes over larger ones (**Fig. 6**). For both p53TAD and Pup, clusters with 2 or 3 residues make up more than 50% of all clusters and clusters with more than 10 residues have notably low occurrence, although their formation could be functionally relevant during molecular recognition events. Because clusters consisting of residue pairs dominate intra-residual interactions in both IDPs, further analysis of the interaction network was performed based on pairwise contacts. Contact maps were generated for p53TAD and Pup averaged over all MD trajectories and pairwise contacts that have occupancies larger than 0.2 visualized as separate graphs (**Fig. 4E,F**). Instantaneous clusters can belong to such larger graphs as exemplified by clusters A1.1, A1.2, A1.3 for p53TAD and clusters B1.1 and B1.2 for Pup (**Fig. 4E,F**). The dominant clusters are characterized by a mix of hydrogen bonds, salt bridges (e.g., involving Arg65 in cluster A2, Arg8 in star-like cluster B1.2, and Arg56 in cluster B3), hydrophobic and aromatic interactions (e.g., Phe19, Leu22/25/26, and Trp23 in cluster A1). These are consistent with the driving forces attributed to liquid-liquid phase separation, namely intermolecular contacts among aromatic residues,(74–76) electrostatic interactions,(77–79) and hydrophobic interactions. (80)

The majority of clusters are linear graphs with few circular sub-graphs leading to the linear relationship between the number of nodes and number of edges (**Fig. 6B**). Acidic residues tend to have low cluster participation whereas arginine residues have the highest participation in both proteins (**Fig. 5A,B**). This difference in cluster participation between cationic and anionic residues is also evident in **Fig. 5C,D**. Among the neutral amino acids, those with larger side-chains are more prone to interactions with non-neighboring residues due to their intrinsically larger distance range. In fact, Pro, Val, Ser, Ala, Gly have the lowest interaction propensities among neutral residues and among pairs of chemically similar residues, such as Gln vs. Asn and Leu vs. Val, the larger residue (Gln, Leu) dominates the smaller one (Asn, Val).

A primary biological function of p53TAD is to negatively regulate p53 by interacting with the ubiquitin ligases MDM2 and MDMX for the degradation of p53. This interaction is one of the earliest and best studied interactions between an IDP and a folded protein both by experiment(65, 66, 81) and computation.(82) In order to better understand the molecular recognition mechanism underlying the formation of this complex, a realistic and accurate description of the free state of p53TAD is of central importance. For MD studies, the choice of the protocol, especially of the force field and water model, is consequential. A recent unbiased REMD study of free p53TAD reported the detailed comparison using five different MD force fields all without residue-specific backbone potentials. Based on 1-μs long replicas major differences were revealed in terms of the structural propensities among them and also with respect to experimental data.(83) An even longer simulation of residues 10–39 of p53TAD for a total length of 1.4 ms analyzed by Markov state models identified substantial populations of β-sheets across the sequence,(84) a behavior that is at variance with the above mentioned REMD ensembles(83) as well as with experimental solution NMR data.(65) These together with many other studies show that force fields need to be chosen following extensive testing to ensure that long trajectories, generated with considerable computational effort, offer the most realistic biophysical insights about these highly complex, heterogeneous systems.

In addition to forming transient intramolecular contacts, IDPs can also dynamically interact with other IDPs driving the formation of liquid-liquid phase separation. With a rapidly increasing body of experimental data on LLPS condensates,(9, 10, 85) all-atom MD simulations have an important role to play for a mechanistic understanding of emerging phase separation properties. Since the molecular driving forces of LLPS are the same as for intramolecular IDP interactions,(86) such as those described here, the optimal accuracy of force fields along with adequate sampling schemes of the heterogeneous condensate environment will be key for the quantitative interpretation of experimental data, allowing the prediction of condensate formation and eventually may open the way for new interventional approaches to actively reprogram condensates and their properties.

Although a possible role of Pup in LLPS is not known, LLPS involving full-length p53 has been documented and p53TAD has been implicated in both phase separation and oncogenic amyloid aggregation.(87, 88) Multivalent electrostatic interactions between the N-terminal domain, p53TAD, and the C-terminal domain were identified as critical for LLPS, which were shown to be positively modulated through molecular crowding and negatively modulated by the addition of DNA and ATP molecules and post-translational modification. It was suggested that compartmentalization of p53 into the droplets suppresses its transcriptional regulatory function, while its release from droplets under cellular stress can activate p53.(87) These findings point to the need for the comprehensive characterization of these intermolecular interactions at residue- and atomic-level resolution. The agreement with experiment reported here clearly suggests that MD methodology has reached a level of accuracy allowing it to make critical contributions toward this goal.

The results of our study further advance the long-held premise of MD simulations to realistically describe IDP ensembles on their native dynamics time scales toward the better understanding of their biophysical properties and biological function. For the two IDPs p53TAD and Pup, the use of REMD allows the adequate sampling of conformational space for the generation of a representative set of initial structures that are then subjected to long, continuous MD simulations. The close agreement found for the extendedness of the simulated IDPs with experiment and polymer theory suggests an appropriate balance between the ff99SBnmr2 force field and the TIP4P-D water model at the global scale. It favorably complements the authentic IDP behavior achieved by this protocol on the local scale in terms of its compliance at the individual residue level with coil libraries, scalar couplings, and chemical shifts. In addition to the realistic modeling of ensemble properties, our protocol also reproduces motional amplitudes and time scales encoded in quantitative NMR spin relaxation data with near experimental accuracy suggesting that the dominant minima of the free energy surface together with their many low-lying transition states are realistically captured by this comprehensive computational framework. These results prompted a more detailed analysis of short-lived inter-residue interactions, which was achieved by graph theory revealing characteristic inter-residue contact patterns and the extraction of residue-type specific interaction propensities. The realistic IDP conformational dynamics model achieved by the protocol described here advances our increasingly mechanistic and predictive understanding of IDPs along with their interactions and binding properties with ordered and disordered molecular targets ranging from regulatory pathways to emerging LLPS phenomena.

## METHODS

### Molecular dynamics simulations

Fully extended structures of p53TAD and Pup were prepared using the LEaP program in AmberTools16.(89) After equilibration, they were used to run replica-exchange MD (REMD) simulations for the sampling of conformational space (36 replicas for each IDP covering a temperature range from 298–353 K for p53TAD and 298–365 K for Pup, see Supplementary Material) with each replica being 1 μs of length. Exchange was attempted every 10 ps and the exchange probability was about 0.3. For each IDP, 10 structures were randomly selected from the room-temperature (298 K) REMD ensemble and used as initial structures to run free MD simulations for 1 μs in the NPT ensemble at 300 K and 1 atm. The protein force field and water model used in all simulations were AMBER ff99SBnmr2 and TIP4P-D.

All MD simulations were performed using the GROMACS 2020.2 package.(90) The integration time step was set to 2 fs with all bond lengths containing hydrogen atoms constrained by the LINCS algorithm. Na^+^ or Cl^-^ ions were added to neutralize the total charge of the system. A 10 Å cutoff was used for all van der Waals and electrostatic interactions. Particle-mesh Ewald summation with a grid spacing of 1.2 Å was used to calculate long-range electrostatic interactions. A cubic simulation box extending 8 Å from the protein surface in all three dimensions was used. Energy minimization was performed using the steepest descent algorithm for 50,000 steps. The system was simulated for 100 ps at constant temperature and constant volume with all protein heavy atoms positionally fixed. The pressure was then coupled to 1 atm and the system was simulated for another 100 ps. The final production run of 1 μs length was performed in the NPT ensemble at 300 K and 1 atm. For simulation details, see **Table S1**.

### Radius of gyration tensor calculations and derived quantities

In order to map the global shape of p53TAD and Pup conformers, radius of gyration tensors were computed as 3×3 matrices ***S*** from each snapshot of the room-temperature REMD ensemble and the free MD simulations as follows:(91)

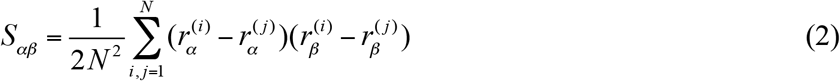

where 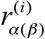 is cartesian coordinate α (β) (= *x*, *y*, *z*) of atom *i* in the coordinate system that has its origin in the center of mass of the molecule. Diagonalization of ***S*** yields three non-negative eigenvalues 0 ≤ *λ*_1_ ≤ *λ*_2_ ≤ *λ*_3_ from which the *radius of gyration R_g_* is obtained, *R_g_* = (*λ*_1_ + *λ*_2_ + *λ*_3_)^1/2^, the *asphericity A*,(91, 92)

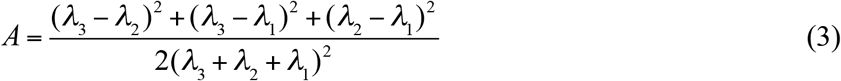

and the *prolateness P*,(93)

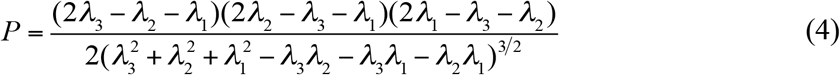

The asphericity measures the degree to which the three axis lengths of the ellipsoid of inertia (eigenvalues) are equal, whereas the prolateness *P* indicates whether the largest or smallest axis length is closer to the middle axis length. *P* takes values between −1 and 1, quantifying the transition from oblate to prolate shapes. Normalized time-correlation functions of *R*_g_(t), made offset-free, were computed according to

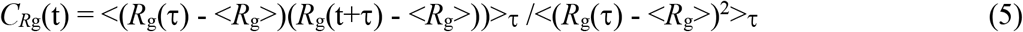

as an average over all 1-μs MD trajectories.

According to polymer theory, for an unfolded polymer the ensemble-averaged *R*_g_ scales with the number of residues *N* as(61)

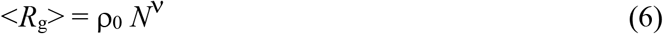

where ρ_0_ is a constant reflecting the average size of a residue and the Flory exponent v determines the overall compactness of the polymer serving as a reference.

### Back-calculation of *R*_1_, *R*_2_ relaxation rates

For IDPs, the normalized time-autocorrelation function *C*(*t*) of the lattice part of the spin-relaxation active magnetic dipole-dipole interaction cannot be factorized into an overall tumbling part and an internal dynamics part. Rather, we compute the full *C*(*t*) directly from an MD trajectory using the second-order Legendre polynomial:

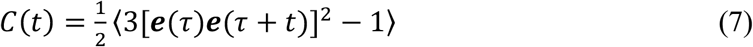

where ***e***(*t*) is the unit vector defining the ^15^N–^1^H bond orientation whereby snapshots were *not* aligned with respect to a reference snapshot. The angular brackets indicate averaging from time τ = 0 to *T*_MD_ – *t*, where *T*_MD_ is the total trajectory length. The calculation of *C*(*t*) was efficiently performed by the fast Fourier transform (FFT) using the Wiener–Khinchin theorem. For acceptable statistical convergence, the analysis of *C*(*t*) was limited to its initial portion from t = 0 - *T*_MD_ /3. Next, a multiexpoential decay function was fitted to *C*(*t*):(94)

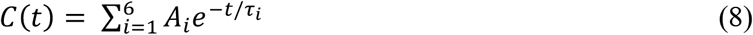

where *A_i_* and *τ_i_* are the best fitting parameters subject to the conditions:

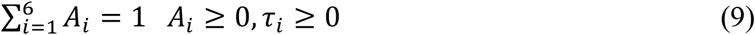

The spectral density function *J*(ω) can be then analytically obtained via Fourier transformation of *C*(*t*):

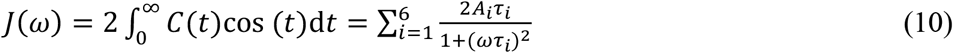

NMR spin relaxation parameters *R*_1_ and *R*_2_ were then computed using the standard expressions:(95–98)

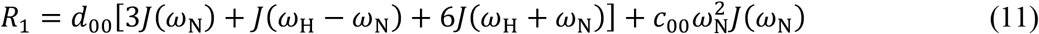

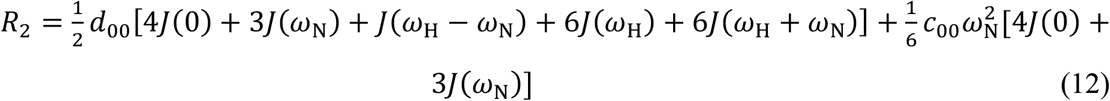

where 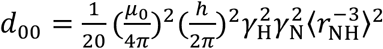 and 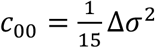. μ_0_ is the permeability of vacuum, *h* is Plank’s constant, γ_H_ and γ_N_ are the gyromagnetic ratios of ^1^H and ^15^N, and *r*_NH_ = 1.02 Å is the backbone N-H bond length. The ^15^N chemical shift anisotropy was set to Δσ = −160 ppm.

### Analysis of inter-residue contacts and residue clusters by graph theory

Contact analysis was performed on all snapshots of the MD simulations of both p53TAD and Pup. A contact is considered formed when the nearest distance between atoms from two different residues is smaller than 4 Å. First-neighbor contacts (between residues *i*,*i*+1), and second-neighbor contacts (between residues *i*,*i*+2) were excluded since they are present for most residues. For each residue in p53TAD and Pup, the total number of contacts formed by a particular residue is determined and normalized by the number of MD snapshots. Each snapshot was converted to a graph where residues are represented as nodes and contacts between two residues are represented as edges between them. The initial graph was then decomposed into a maximal number of disconnected graph components called *clusters*, i.e. there is no edge between any node in the cluster and any node outside the cluster. The size of a cluster corresponds to the number of its nodes.

## ACKNOWLEDGEMENTS

We thank Dr. Da-Wei Li for helping with the graph theoretical analysis. MD and REMD simulations were performed at the Ohio Supercomputer Center. The authors declare that they have no competing interests. All data needed to evaluate the conclusions in the paper are present in the paper and/or the Supporting Information.

## SUPPORTING INFORMATION

Fig. S1. Radius of gyration of the IDPs p53TAD and Pup in 10 1-μs MD trajectories each at 300 K with starting structures randomly chosen from replica exchange simulations.

Fig. S2. Mean *R*_1_, *R*_2_ errors from 10 1-μs MD simulations of p53TAD and Pup in comparison with experiment.

Fig. S3. Back-calculated *R*_1_, *R*_2_ ^15^N backbone spin relaxation rates from microsecond MD simulations of p53TAD and Pup excluding atypical trajectories in comparison with experiment.

Fig. S4. Comparisons of experimental and predicted chemical shifts of p53TAD.

Fig. S5. Experimental and MD-derived secondary structure propensities of p53TAD.

Fig. S6. Average number of contacts formed by a particular residue in p53TAD and Pup per snapshot using only side-chain atoms.

Fig. S7. Contact propensities according to amino-acid residue type for both proteins combined.

Table S1. MD and REMD simulation details for p53TAD and Pup.

Table S2. Most frequent pairwise residue contacts in p53TAD from MD simulations.

Table S3. Most frequent pairwise residue contacts in Pup from MD simulations.

## Notes

### Competing Interest Statement

The authors have declared no competing interest.

